# BindSpace: decoding transcription factor binding signals by large-scale joint embedding

**DOI:** 10.1101/359539

**Authors:** Han Yuan, Meghana Kshirsagar, Lee Zamparo, Yuheng Lu, Christina S. Leslie

## Abstract

Decoding transcription factor (TF) binding signals in genomic DNA is a fundamental problem. Here we present a prediction model called BindSpace that learns to embed DNA sequences and TF class/family labels into the same space. By training on binding data for hundreds of TFs and embedding over 1M DNA sequences, BindSpace achieves state-of-the-art multiclass binding prediction performance, *in vitro* and *in vivo*, and can distinguish signals of closely related TFs.

## Main

Direct measurement of genome-wide transcription factor (TF) occupancy for all expressed factors in a cell type of interest is practically infeasible outside of large consortium projects. Therefore, computational prediction of TF binding to cognate sites at relevant loci – e.g. chromatin accessible regions or putative enhancers defined by active histone marks – is of critical importance. Massive efforts to define the intrinsic binding affinities of TFs by protein binding microarray (PBMs)^1^, cognate site identification (CSI)^2^, genomic-context PBM (gc-PBM)^3^, mechanically induced trapping of molecular interactions (MITOMI)^4^ and high-throughput SELEX followed by sequencing (HT-SELEX)^5^ provide large-scale data sets for training binding models. However, these *in vitro* binding experiments are typically summarized as position-specific weight matrix (PWM) motifs, losing both specificity and sensitivity and leading to near-identical motifs for closely related TFs. Supervised learning methods have improved accuracy of discrimination between bound and unbound sequences of individual TFs^6–9^ but have not addressed the multiclass nature of the problem and therefore are not optimized to distinguish between TFs with similar binding signals.

Here we present a novel multiclass and multilabel method to jointly learn binding preferences of hundreds of assayed TFs by embedding their bound/unbound DNA sequences and class labels into a common space. Our method, called BindSpace, learns accurate binding models for individual TFs while enabling discrimination between different TFs in the same family. To train BindSpace, we combined HT-SELEX *in vitro* binding experiments for 461 mouse and human TFs from previous large-scale studies^5,8^. After applying rigorous quality control (**Methods**), we used 270 experiments for 243 transcription factors for our training set. The top 2000 enriched probes from each of these experiments were used as positive examples, yielding over 500K positive training sequences. We randomly sampled universal negatives from initial HT-SELEX probe libraries as well as non-accessible genomic regions to obtain ~500K negative training sequences (**Methods**). Each sequence is represented as a bag of 8-mers, each containing up to two consecutive wild cards, and each bag is associated either with both a TF label (e.g. HOXA2) and a TF family label (e.g. Homeodomain) or with a universal negative label. In this study, we used two thirds of the HT-SELEX data for training and one third for testing, and we performed 5-fold cross validation on the training data for hyperparameter tuning (**Methods**).

BindSpace is a multiclass and multilabel supervised embedding model that learns to map *k*-mer features, sequence examples, and TF/family labels to a shared high dimensional space. We adapted a general purpose embedding framework from natural language processing (NLP) called StarSpace^10^ to train our model (**Methods**). In NLP applications, StarSpace learns an embedding of words into a semantic embedding space, which defines an embedding of sentences or documents; here, *k*-mers are analogous to words and DNA probe sequences to sentences. BindSpace maximizes the similarity between a sequence example and its corresponding labels in the embedding space, while minimizing the similarity between the example and other labels. The label of a new sequence is determined by the similarity of its embedded vector to all the labels in the embedding space (**Fig. 1**). In computational biology, the idea of mapping sequences into a semantic embedding space was previously used for protein domain sequences for the task of remote homology detection^11^.

**Figure 1.**
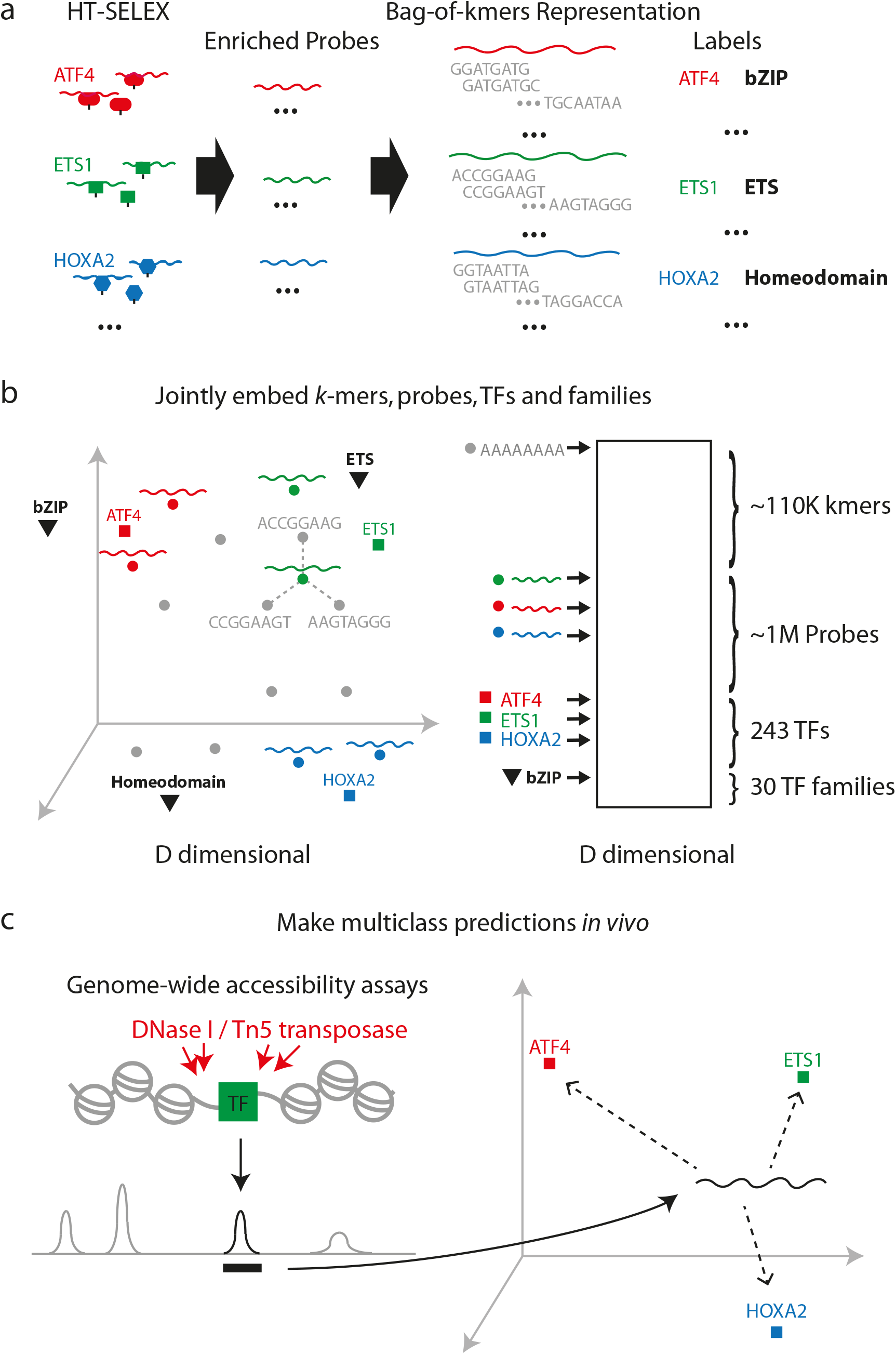
A schematic overview of BindSpace. BindSpace is an embedding approach that jointly learns binding models for hundreds of TFs. **(a)** Enriched DNA probes for TF HT-SELEX experiments are represented as bags of *k*-mers, namely 8-mers with up to two consecutive wildcards. **(b)** BindSpace learns an embedding of *k*-mers (grey dots), DNA probes (colored dots), TF class labels (colored squares), and TF family labels (black triangles) into the same high-dimensional space. The embedding of a probe sequence is specified by the embedding of its constituent *k*-mers. Unbound HT-SELEX probes and inaccessible genomic sequences serve as universal negative examples. **(c)** The trained BindSpace model can be used for multiclass prediction of TF binding *in vitro* or *in vivo*.

Intuitively with BindSpace, we learn a model where TFs with similar binding specificity are close to each other and their training examples in the embedding space, while TFs with very different binding specificity are farther away. By visualizing the embedding space in 2D using t-distributed stochastic neighbor embedding (t-SNE) (details in **Methods**), we can view the relative embedding of probes, TF labels, and family labels (**Fig. 2a**). Probe sequences clearly cluster in the embedding space, with TF families occupying their own territories (labels and probes for 13 major shown; full visualization of all TF labels in **Supplementary Fig. 1**). Zooming into a particular TF family such as bZIP shows that the model also learns family substructure (**Supplementary Fig. 2**). For example, human and mouse orthologs are embedded at very similar positions (Jdp2 and JDP2, HLF and Hlf, Dbp and DBP), and the C/EBP TFs similarly embed close to each other.

**Figure 2.**
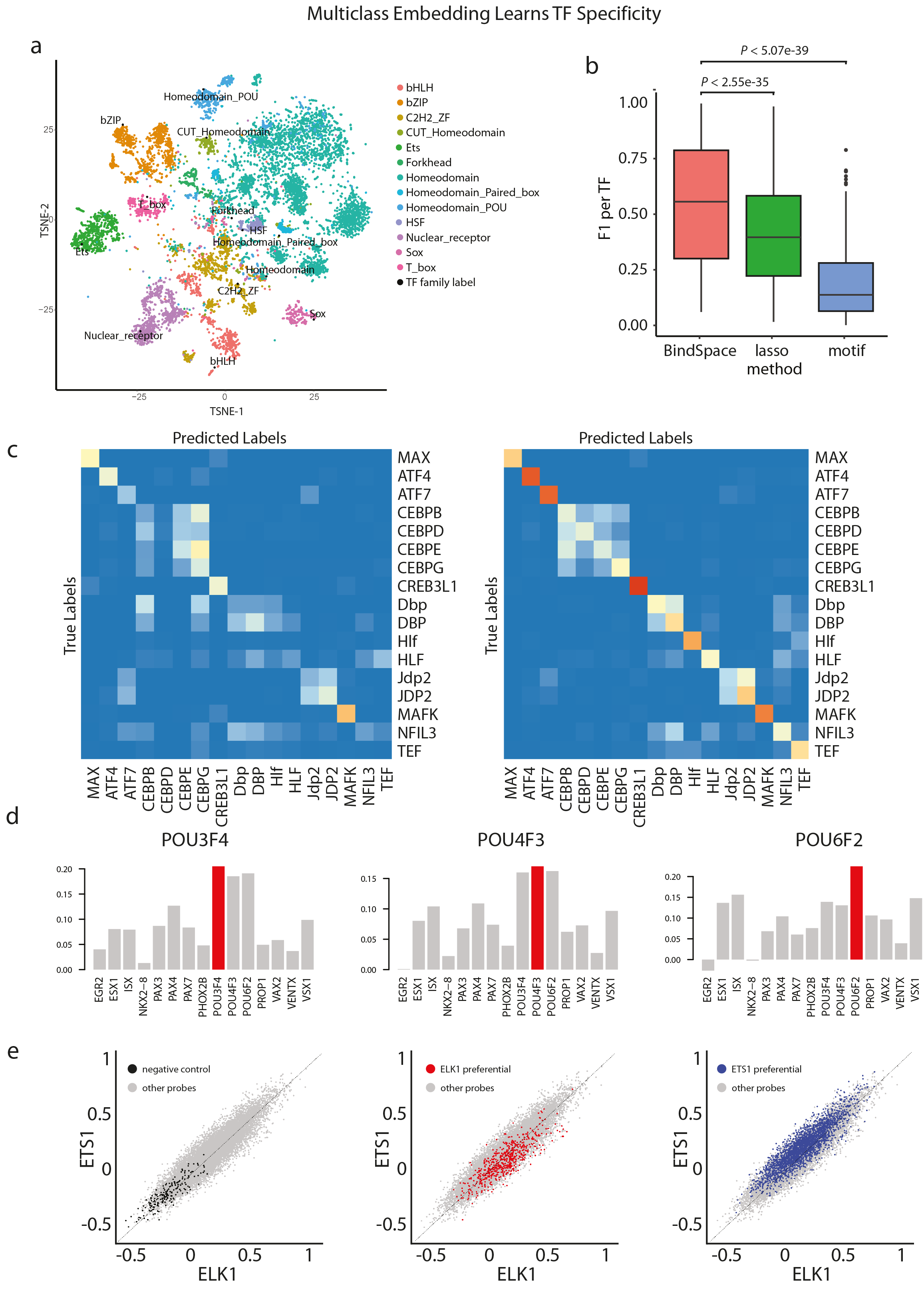
BindSpace accurately predicts TF binding and distinguishes between TF family members *in vitro*. **(a)** t-SNE visualization of the embedding space. Colored points represent embedding of enriched probes from HT-SELEX experiments. Black points represent embedding of TF family labels for 13 major families (families with more than three members). A full t-SNE visualization of all TF labels as well as TF family labels is shown in **Supplementary Fig. 1**. **(b)** Multiclass prediction performance of BindSpace, motif and lasso one-versus-all methods on held-out HT-SELEX probes. Classification performance is measured by F1 score for each TF class. **(c)** Multiclass prediction performance of motif scoring (left) and BindSpace (right) as shown by confusion matrices normalized by class support for the bZIP family. Rows are true labels, and columns are predicted labels. **(d)** For each of the 15 PBM experiments that overlap between BAR15A and BindSpace, we predicted the binding affinity of TFs to the top intensity probes from each of these PBM experiments using BindSpace. Here from left to right we show the results for three PBM experiments of paralogous TFs POU3F4, POU4F3 and POU6F2. In each plot, we show the correlations of estimated affinities to measured PBM intensities. **(e)** For all three plots, we show on the x-axis the BindSpace predicted affinity of all gcPBM probes to ELK1, and on the y-axis the BindSpace predicted affinity to ETS1. Negative control probes (those that do not bind either ELK1 or ETS1 *in vivo*) are labeled in black (left); high affinity ELK1 preferential probes are labeled in red (middle), and high affinity ETS1 preferential probes are labeled in blue (right).

We first evaluated our model on the held-out HT-SELEX test data and compared against FIMO motif scoring using published HT-SELEX PWM motifs curated using a semi-automated algorithm^5,12^; note that no probe data was held out for estimating these PWMs, thereby giving the motif-based method an unfair advantage. Nonetheless, BindSpace significantly outperforms motif-based prediction on the multiclass classification task, presumably because motifs are fit individually rather than in a multiclass manner (*P* < 5.07e-39, signed rank test, **Fig. 2b**). BindSpace also outperforms one-versus-all LASSO classifiers trained using 8-mers with up to two consecutive wild cards as features (*P* < 2.55e-35, signed rank test, **Fig. 2b**). As previously shown, such sparse linear models with *k*-mer features offer competitive performance with state-of-the-art methods to model TF binding specificity^7^. Comparing the confusion matrices within each TF family obtained by BindSpace versus motif scoring, the multiclass performance advantage of BindSpace is apparent (**Fig. 2c**, **Supplementary Fig. 3, 4**). We also observe that some paralogs are much harder to distinguish than the rest of the family, such as the C/EBP subfamily (**Fig. 2c**).

We then evaluated BindSpace on independent PBM data sets to confirm that we could reproducibly distinguish TFs within the same family across *in vitro* platforms. We downloaded raw PBM intensity data for the BAR15A data set^13^, which contains PBM data for 41 TFs, 15 of which overlap with those in BindSpace (1 in the C2H2_ZF family, 14 in the homeodomain family). We predicted the binding affinity of all 15 TFs to the top intensity probes from each of these PBM experiments using BindSpace (**Methods**). For 9 of the 15 TFs, the PBM intensity correlated best with the BindSpace model for the same TF (**Supplementary Fig. 5**), even for paralogs such as POU3F4, POU4F3 and POU6F2 (**Fig. 2d**). For PAX family paralogs, BindSpace successfully distinguished PAX4 and PAX7 from the rest but had more difficulty with PAX3.

Shen *et al.* recently described custom gcPBM experiments and a computational framework to detect subtle differences in binding preferences between paralogous TF pairs^9^. Specifically, for the paralogous pair Ets1 and Elk1 (the only pair that overlaps with BindSpace), they designed a custom gcPBM array of 13,765 probes and annotated each probe as Elk1 preferential, Ets1 preferential or neither. Using BindSpace, we predicted the affinity of gcPBM probes to ELK1 and ETS1 (**Methods**). We found that high affinity probes annotated as Elk1 preferential by Shen *et al.* also have higher BindSpace scores for ELK1 (*P*< 3.82e-18, signed rank test), and high affinity Ets1 preferential probes have higher BindSpace scores for ETS1 (*P* < 1e-64, signed rank test) (**Fig. 2e, Supplementary Fig. 6**). This suggests that BindSpace can capture subtle differences in binding between paralogous TFs. In addition, when we focused on probes including specific core sequences (GGAT, GGAAC and CAGGAA), we recapitulated reported paralog specific recognition: probes containing GGAT/ATCC and CAGGAA/TTCCTG were scored significantly higher by the BindSpace ETS1 model (*P* < 1e-64, *P* < 1e-64, signed rank test), and probes containing GGAAC/GTTCC were scored significantly higher by ELK1 model (*P* < 1.14e-14, signed rank test), consistent with gcPBM measurements (**Supplementary Fig. 6**).

BindSpace also accurately identified TF binding sites and distinguished the binding specificity of related TFs *in vivo*. We first evaluated BindSpace on the task of distinguishing TF binding sites from flanking sequences, using ENCODE ChIP-seq for 39 TFs represented in the model. We used both BindSpace and FIMO motif scoring to distinguish the top 5,000 peaks from flanking regions (**Methods**). BindSpace significantly outperformed motif scoring on this binary recognition task as measured by F1 score (*P* < 3.75e-4, signed rank test, **Fig. 3a**, **Supplementary Table 2**). Scoring every 20bp window of the ChIP-seq peak for TF affinity using BindSpace, we observed that many bZIP transcription factors (especially MAFK and CEBPB) have binding sites well aligned at the center of ChIP-seq summits, whereas most other TF have binding signals widely distributed over ChIP peaks (**Supplementary Fig. 7**).

**Figure 3.**
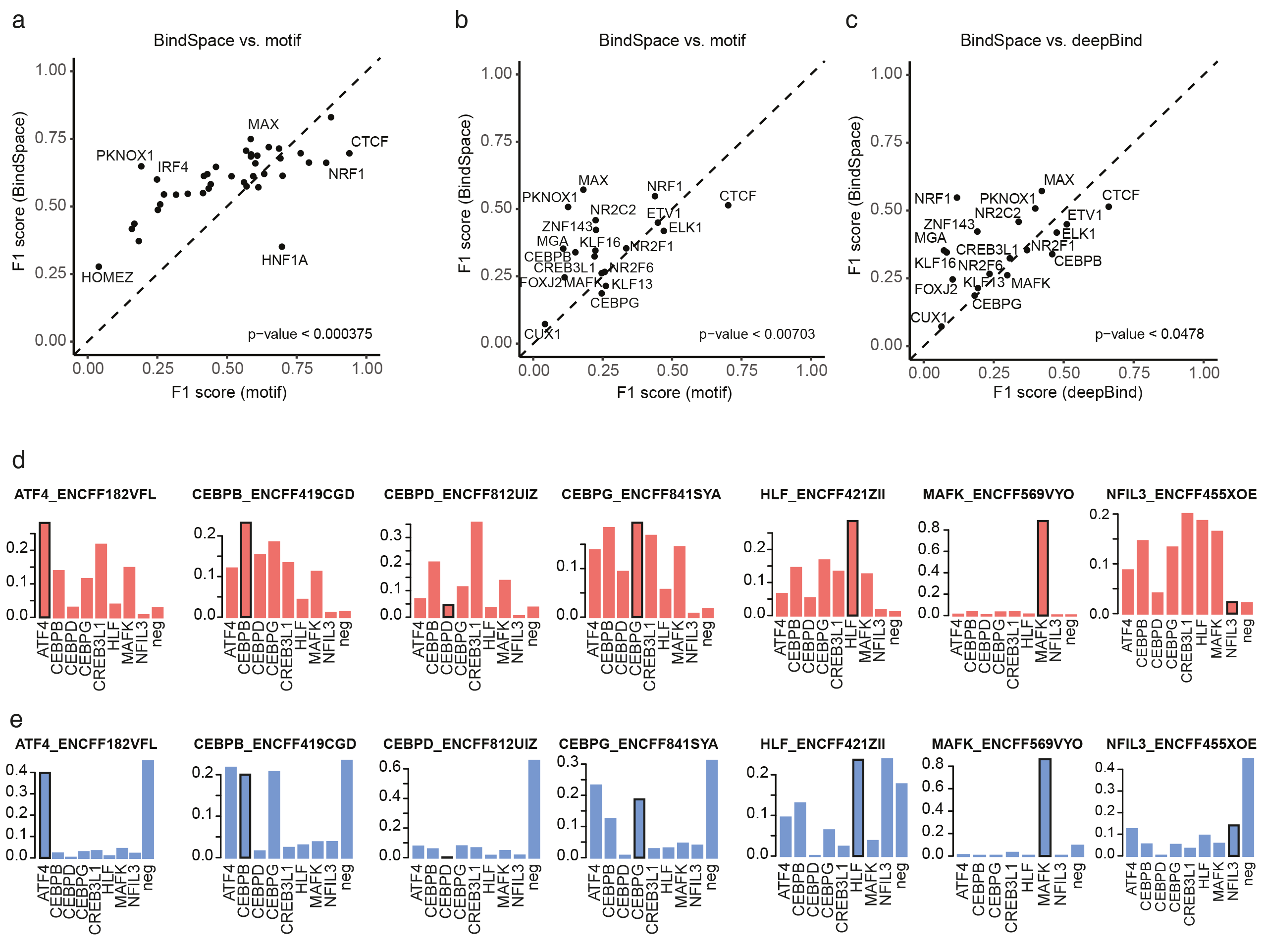
BindSpace predicts binding of TFs and distinguishes between paralogous TF binding sites *in vivo*. **(a)** Performance of BindSpace versus motif scoring as measured by F1 score on the task of distinguishing TF ChIP-seq peaks from flanking sequences. **(b)** Performance of BindSpace versus motif method as measured by F1 score on the binary task of distinguish TF-associated ATAC-seq peaks versus TF-free ATAC-seq peaks in K562. **(c)** Performance of BindSpace versus DeepBind on the binary task of distinguishing TF-bound ATAC-seq peaks from unbound ATAC-seq peaks in K562. **(d)** Multiclass classification performance of BindSpace for bZIP family members in HepG2 cell line. For each plot, we performed multiclass classification using BindSpace on the top 10,000 peaks for bZIP TF ChIP-seq and show the proportion of predicted labels for each model. BindSpace successfully ranked the ChIP-ed TF above other family members in 5 out of 7 cases. **(e)** As in (d), reporting classification performance for motif scoring instead of BindSpace. Motif scoring only ranks the ChIP-ed TF above other family members in 3 out of 7 cases.

We next evaluated performance for the more practical task of predicting TF binding versus non-binding at chromatin accessible regions in a given cell type. We processed publicly available ATAC-seq data for K562 and GM12878 (**Methods**) and downloaded available ENCODE ChIP-seq data for these cell lines. Restricting to TFs represented in BindSpace, we found 19 TFs in K562 and 11 TFs in GM12878 with ChIP-seq data. For K562, BindSpace significantly outperformed motifs in discriminating ATAC-seq peaks with occupancy for a given TF versus unbound peaks as measured by F1 score (*P* < 0.00703, signed rank test, **Fig. 3b**, **Supplementary Table 3**). BindSpace also significantly outperformed DeepBind, a deep learning method trained on HT-SELEX data set, on this task (*P* < 0.0478, signed rank test, **Fig. 3c**)^14^. BindSpace had similar performance advantages on the GM12878 data set (*P* < 0.0451, signed rank test, **Supplementary Fig. 8**, **Supplementary Table 4**).

Finally, we tested BindSpace’s ability to distinguish between the binding of TFs in the same family *in vivo*. We searched ENCODE ChIP-seq experiments in major cell lines for TFs that overlap those in BindSpace and found 115 such ChIP experiments. For each of these, we performed multiclass classification on the top 10,000 peaks to test if we could distinguish binding of the ChIP-ed TF from TFs in the same family. For some TF families, BindSpace made much better multiclass predictions than motif scoring. In particular, the bZIP family has the largest number of members represented both in ENCODE ChIP-seq and BindSpace (22 ChIP-seq experiments for 8 TFs, and 7 TFs ChIP-ed in the HepG2 cell line). Examining BindSpace predictions for bZIP ChIP-seq in HepG2, BindSpace successfully ranked the ChIP-ed TF above other family members in 5 out of 7 cases (**Fig. 3d**; predictions for all 22 bZIP ChIP-seq experiments **see Supplementary Fig. 9**). Motif scoring, on the other hand, only predicted the correct TF for 3 out of 7 cases. Motif scoring often gets confused between paralogs. For example, the most enriched motif in CEBPB ChIP-seq is ATF4, and then CEBPG; the most enriched motif in CEBPG ChIP-seq is also ATF4; and the most enriched motif in HLF ChIP-seq is NFIL3 (**Fig. 3e**).

BindSpace is a powerful and scalable machine learning approach to leverage massive *in vitro* binding data sets to decode TF binding signals in DNA sequences. By solving the underlying “many-class” classification problem through a joint embedding space for sequences and TF class labels, BindSpace is often able to distinguish between the binding preferences of closely related TFs, both *in vitro* and *in vivo*. While we have trained here on HT-SELEX experiments, we envision extensions where we train across multiple *in vitro* platforms to expand the repertoire of TFs and families represented in the model. Moreover, we expect that the BindSpace embedding will provide a useful feature representation on which to build more sophisticated models, such as deep learning models, for diverse problems in genomics. Open source code for BindSpace as well as the trained BindSpace model are freely available for download (https://bitbucket.org/hy395/selex_embed for source code of training; https://bitbucket.org/hy395/bindspace for R-package for making prediction with trained models).

## Supplementary Figure Legends

### Supplementary Figure 1. t-SNE visualization of the embedding space

Colored points represent embedding of enriched probes from HT-SELEX experiments. Black points and labels represent embedding of TFs (except for Homeodomain TFs). Homeodomain TFs are shown by gray points, and the associated labels are not shown because there are too many of them. Red points and labels represent embedding of 13 major families (families with more than three members).

### Supplementary Figure 2. t-SNE visualization of the embedding space for the bZIP family

The same t-SNE visualization as Supplementary Fig. 1, but zooming into the top left corner for a close look at the bZIP family. Colored points represent embedding of enriched probes from bZIP HT-SELEX experiments. Black points and labels represent embedding of TFs. Red point and label represent embedding of bZIP family label.

### Supplementary Figure 3. Confusion matrices for BindSpace and motif scoring

Multiclass prediction performance for motif scoring (left) and BindSpace (right) as shown by confusion matrices normalized by class support for all 13 major families. Rows are true labels, and columns are predicted labels.

### Supplementary Figure 4. Confusion matrices for BindSpace and motif scoring (continued)

Multiclass prediction performance for motif scoring (left) and BindSpace (right) as shown by confusion matrices normalized by class support for all 13 major families. Rows are true labels, and columns are predicted labels.

### Supplementary Figure 5. BindSpace analysis of PBM data

For each of the 15 PBM experiments that overlap between BAR15A and BindSpace, we predicted the binding affinity of TFs to the top intensity probes from each PBM experiment using BindSpace. In each plot, we show the correlation of estimated affinities to measured PBM intensities. For 9 of 15 experiments, the PBM intensity correlated best with the BindSpace model for the assayed TF.

### Supplementary Figure 6. BindSpace analysis of gcPBM data

**(a)** Reported intensity of Ets1 and Elk1 to gcPBM probes as designed and measured by Shen *et al.* From left to right, we labeled the negative control probes in gray (left), high affinity Elk1 preferential probes in red (middle), and high affinity Ets1 preferential probes in blue. The same set of negative control probes, high affinity Elk1 preferential probes and high affinity Ets1 preferential probes were for our analysis in **Fig. 2e**.

**(b)**BindSpace predicted affinity of all gcPBM probes to ELK1 and ETS1. From left to right, we labeled probes containing GGAT/ATCC in blue, GGAAC/GTTCC in red and CAGGAA/TTCCTG in blue.

### Supplementary Figure 7. Spatial analysis of BindSpace scores on ChIP-seq peaks

BindSpace scores for ChIP-seq peaks centered around summit at 20bp resolution. TFs belonging to the same family are surrounded by a red box.

### Supplementary Figure 8. Binary prediction performance of TF binding at ATAC-seq peaks

Performance of BindSpace versus motif scoring on the binary task of distinguish TF-associated ATAC-seq peaks versus TF-free ATAC-seq peaks in GM12878.

### Supplementary Figure 9. Multiclass prediction performance of TF binding at bZIP ChIP-seq peaks

Multiclass classification performance of BindSpace and motif scoring for 22 ENCODE ChIP-seq experiments for bZIP family members. For each plot, we performed multiclass classification using BindSpace on the top 10,000 peaks for bZIP TF ChIP-seq and show the proportion of predicted labels for each model. Red barplots show BindSpace predictions and blue barplots show motif prediction. The TF being ChIP-ed in each experiment is labeled with a black box around the bar. In all 22 experiments, BindSpace successfully ranked the ChIP-ed TF above other family members in 16 out of 22 cases, as compared to 8 out of 22 cases for motif scoring.

### Supplementary Figure 10. HT-SELEX quality control

Quality of raw HT-SELEX data. For each HT-SELEX experiment, we computed the Spearman correlation of probe enrichment scores between cycle 3 and cycle 4. This figure shows the distribution of correlations for all experiments. We filtered out experiments with a correlation less than 0.8.

### Supplementary Figure 11. Results of HT-SELEX quality filtering

Our filtering steps improved overall HT-SELEX data quality. We computed the consistency between replicate HT-SELEX experiments or experiments on orthologous (mouse/human) TFs. We defined the consistency between experiment A and B as the percentage overlap between the top 100 enriched 8-mers in the two experiments. Here we show the consistency between all pairs before quality control, and consistency between all remaining pairs after quality control.

### Supplementary Figure 12. BindSpace hyperparameter tuning

This figure shows the changes in BindSpace mean F1 scores in cross-validation with respect to changes in the hyperparamters: learning rate, dimension, number of negative samples and dropout rate.

## Supplementary Tables

**Supplementary Table 1**. Information of 270 HT-SELEX experiments used for training, and also the number of unique probes and non-unique probes among top 2000 in each experiment.

**Supplementary Table 2**. Performance of BindSpace and motif to distinguish ChIP-seq peaks versus flanking sequences.

**Supplementary Table 3**. Performance of BindSpace, motif scoring, and DeepBind to distinguish TF-bound ATAC-seq peaks versus unbound ATAC-seq peaks in K562.

**Supplementary Table 4**. Performance of BindSpace and motif scoring to distinguish TF-bound ATAC-seq peaks versus unbound ATAC-seq peaks in GM12878.

## Methods

### HT-SELEX quality control

We used public HT-SELEX data sets to train BindSpace and one-vs-all lasso TF binding specificity models. We combined the HT-SELEX data sequenced in 2013 (ENA accession: ERP001824) and the 2017 re-sequenced libraries (ENA accession: ERP016411), which together include 547 experiments for 461 human or mouse TFs^5,8^.

To perform quality control, we computed an enrichment score for every 8mer in probes selected at each cycle of an experiment relative to the initial library (cycle 0), as in a previous study^8^. We first estimated the frequency of every 8-mer in cycle 0 using a fifth-order Markov model^15^. Then, for cycle i, we computed the enrichment score for each 8-mer as the i-th root of (frequency in cycle i) / (estimated frequency in cycle 0).

For each experiment, we then performed quality control filtering based on the following procedure:

1. For experiments with 14bp or 20bp probes, cycle 4 was used to derive enriched probes.
2. For experiments with 30bp and 40bp probes, one of cycle 4, 5 or 6 was used to derive enriched probes, because more cycles of selection are required to enrich for longer probes.
3. Experiments were excluded if the Spearman correlation of 8-mer enrichment scores between cycle 3 and cycle 4 was below 0.8. (The correlation distribution between cycle 3 and cycle 4 for all experiments is shown in **Supplementary Fig. 10**).
4. Experiments were excluded if the selected cycle did not have good probe enrichment or diversity (fewer than 500 probes with frequency > 10).
5. For the remaining experiments, we removed low complexity probes with a DUST score < 2, where DUST is the BLAST low-complexity masking algorithm^16^. This is based on our observation that some HT-SELEX experiments have non-specific enrichment for low complexity sequences.
6. For remaining experiments with 14bp or 20bp probes, we selected the top 2000 enriched probes (frequency > 10) based on enrichment score.
7. For remaining experiments with 30bp or 40bp probes, we counted the frequency of unique 20bp sequences, and selected the top 2000 enriched 20bp probes based on enrichment score.
8. Finally, experiments where the most frequent 8-mer in the top 2000 probes occurred less than 100 times were removed. This final filter is applied to remove any experiments where the top probes do not enrich for any consensus binding sites.

After quality control, we ended up with 270 high quality experiments covering 243 TFs. 143 of these experiments overlapped with those selected by quality control in a previous study^8^. To show that our filtering steps improved overall data quality, we measured consistency between replicate experiments or experiments of orthologous TFs. Consistency between experiment A and B was defined as the percentage overlap between the top 100 enriched 8-mers in the respective experiments. We found that after removing low quality experiments and probes, we significantly improved overall data quality (*P* < 0.009, rank sum test, **Supplementary Fig. 11**).

### Data preprocessing

The top 2000 enriched probes from each of the 270 experiments that passed quality control were used as examples to train our model. All non-unique probes were removed, and each of the remaining (unique) probes was associated with a TF label and a TF family label. Because 20bp probes have very high diversity, we only had a very small number of non-unique probes (**Supplementary Table 1**). TF family information was obtained from http://cisbp.ccbr.utoronto.ca. Overall we had a total of 505,194 TF-associated probes.

We then randomly selected 252,597 sequences from probes that only present in initial cycles (random negative sequences), and 252,597 20bp sequences from inaccessible region of the genome in K562 cell lines (genomic negative sequences). After filtering out duplicated sequences, we obtained a total of 505,086 unique negative probes, each of which was associated with a universal negative label.

### BindSpace model

Our final data set had 1,010,280 unique sequences, with half of the probes associated to a TF label and a TF family label, and half of the probes associated with a universal negative label. For training and testing, each sequence was represented by a bag of 8-mers with up to two consecutive wild cards.

We used the StarSpace package for training our BindSpace multiclass/multilabel supervised embedding model^10^. We split the whole data set into 2/3 for training and 1/3 for testing with stratified sampling, i.e., each TF class was split into 2/3 for training and 1/3 for testing. We performed 5-fold cross-validation on training data for hyperparameter tuning, and cross-validation performance was measured by average F1 score for all TFs. We then evaluated the performance on test data and compared against other methods.

Finally, we learned a comprehensive BindSpace model with the same hyperparameter settings using all HT-SELEX data. The visualizations and *in vitro* and *in vivo* validation were performed using this model.

### StarSpace

StarSpace is a very powerful general purpose embedding framework introduced in natural language processing^10^. One of its applications is to address multi-class and multi-label classification problems by embedding all examples and labels in a shared space. In this embedding space, it learns to maximize the similarity between an example and its matching labels, while minimizing the similarity between an example and randomly drawn labels. On each batch update, it minimizes the following loss function:

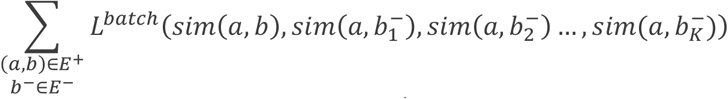

Here (*a, b*) is an example/label pair in the positive set *E*^+^, and 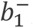, 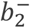…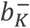 are randomly sampled negative labels from a negative set *E*^−^ generated by *K*-negative sampling^17^. Both the loss function and similarity measure can be considered to be hyperparameters in the model. Optimization is performed by negative sampling and stochastic gradient descent.

### Hyperparameter tuning

The hyperparameters we considered in the StarSpace algorithm included: loss, similarity, learning rate, embedding dimension, maximum number of negatives in a batch update, and dropout rate. Preliminary analysis on a small number of examples (15 classes) showed that hinge loss and dot similarity gave the best performance on DNA sequence data.

We sampled 20 sets of hyperparameters from: learning rate = (0.01, 0.05, 0.1, 0.2), dimension = (50, 100, 150, 200), maxNegSamples = (3, 5, 10, 15), and dropout rate = (0, 0.001, 0.01, 0.1). We compared the cross-validation performance of each of these 20 sets by F1 score for each TF class.

Taking the sampled hyperparameter set with best performance (mean F1 score), we then varied each hyperparameter individually while keeping the rest constant. We measured the performance change with respect to the change in each hyperparameter, again by cross-validation performance on training data measured by F1 score for each TF (**Supplementary Fig. 12**). Based on this analysis, the final hyperparameters that gave the best cross-validation performance were: learning rate = 0.2, dimension = 300, maxNegSamples = 15, dropout rate = 0. We also found that the performance reaches convergence after 50 epochs.

### Making predictions with BindSpace

#### Binary thresholding

BindSpace predicts similarity scores of a new test example to all class labels. Multiclass prediction can be made based on which label has the highest similarity to the test example. However, when we need to make binary prediction for each label, we need to determine a threshold for each label. Therefore, we computed the similarity of each label to all training examples not belonging to this class to generate an empirical null distribution of similarity score. For a new test example, if the similarity score significantly achieves *P* < 0.05 relative to this null distribution, we consider the test example to be positive; otherwise, we consider it to be negative.

#### In vivo evaluation

When evaluating on ChIP-seq or ATAC-seq data, instead of having 20bp probes, we have 150bp or 300bp genomic sequences, respectively. In order to predict on these large genomic regions, we bin into 20bp windows, embed each window into BindSpace, and determine similarity to TF class labels. Then the similarity score of a TF to a given peak is the maximum similarity score over all 20bp windows.

### Motif-based predictions

The motif models we use are from ‘Jolma2013.meme’, downloaded from http://meme-suite.org/db/motifs. They were generated by Jolma *et al.* using a semi-automatic algorithm from the same HT-SELEX data set^5^. We used FIMO with default settings to predict motif hits^12^. For binary prediction problems, we used the default *P* < 1e-4 threshold. For multiclass prediction problems, if none of the motifs satisfied *P* < 1e-4, we predicted the sequence to be a universal negative; otherwise we predicted the TF label based on the most significant motif.

### LASSO one-versus-all multiclass predictions

We used the R package *glmnet* to train a LASSO logistic regression classifier for each TF in a one-versus-all setting in order to make multiclass predictions^18^. During training, for each TF, we considered the 2000 enriched probes as positives and sampled 20,000 sequences from the rest of the training data as negatives. We represented training sequences using 8-mers features with up to 2 consecutive wildcards. We performed feature selection by computing a binomial Z-score for every 8-mer feature based on its enrichment in positive versus negative training examples and chose the top scoring 10,000 8-mers. For testing, we predicted the class of a given test example by assigning it to the TF model that gave highest posterior probability, or assigning it to be a universal negative if all of the models predicted a posterior < 0.5.

### Visualization

We visualized the top 50 probes of each HT-SELEX experiment, TF labels, and TF family labels in a 2D space using an efficient Barnes-Hut implementation of t-SNE^19,20^. To do so, we first computed a similarity matrix of these points (13,500 probes and 274 labels) by dot product. Then we converted this similarity matrix to a dissimilarity matrix and used the R package *Rtsne* with default settings to project these points to 2D, where the dissimilarity is approximated by Euclidean distance^21^.

### BindSpace correlation with PBM data

We downloaded the original PBM intensity data of ‘BAR15A’ from Uniprobe (http://the_brain.bwh.harvard.edu/uniprobe/downloads.php). BAR15A contains PBM data for 41 TFs, 15 of which overlap with transcription factors in BindSpace (1 C2H2_ZF TF, 14 homeodomains). For each TF, we took the top 5000 PBM probes with highest intensity and predicted the binding affinity of all 15 TFs to these 36bp probes using BindSpace. Finally we computed Spearman correlations between true probe intensities with BindSpace predicted intensities for the 15 TFs.

### BindSpace analysis of gcPBM data

We downloaded gcPBM array for Ets1 and Elk1 from GEO (GSE97793). Negative control probes, ELK1 preferential and ETS1 preferential probes were as annotated by Shen *et al*.^9^ In addition, we focused on the group of high affinity ELK1/ETS1 preferential probes in our analysis, i.e. probes that were annotated as ELK1/ETS1 preferential and also had a normalized intensity > 0.7 for either ELK1/ETS1 gcPBM (**Supplementary Fig. 6a**).

### ENCODE ChIP-seq

Optimal IDR thresholded ‘narrowpeak’ files of ENCODE ChIP-seq data were downloaded from the ENCODE portal (https://www.encodeproject.org/) for five major cell lines: GM12878, K562, A549, H1-hESC and HepG2, covering a total of 329 different TFs.

### ATAC-seq data processing

K562 ATAC-seq raw reads were obtained from GSE76224^22^. We combined records SRR3822969 and SRR3822972 to create replicate 1. We combined the other two records to create replicate 2. GM12878 ATAC-seq raw reads were obtained from GSE47753^23^. We used only the samples generated from 50k cells. Because replicate 1 is much more deeply sequenced then rest of the replicates, we combined replicates 2, 3, 4 to obtain replicate 2.

We followed the ENCODE ATAC-seq processing pipeline (https://www.encodeproject.org/atac-seq/) for processing K562 and GM12878 ATAC-seq data. For each of the ATAC-seq samples, we trimmed the raw fastq files of adapters with Trimmomatic with default settings. We then aligned to the hg19 genome using Bowtie2 with default settings. After that we removed duplicate reads and adjusted for Tn5 shifts. Peak calling was performed for each replicate using macs2 with --nomodel --shift -37 --extsize 73. Finally, IDR was performed with the idr package and reproducible peaks were called with an IDR cutoff of 0.05. There were a total of 17,271 reproducible peaks in K562 and a total of 76218 reproducible peaks in GM12878. For efficiency, we only evaluated on the top 10,000 peaks with highest accessibility.

### ChIP-seq peaks versus flanks evaluation

For this task, we used all available ENCODE TF ChIP-seq data in cell lines, which included a total of 329 different TFs, including 39 represented in BindSpace. For each TF covered by BindSpace, we selected a single experiment that gave the most reproducible peaks, selected the top 5000 peaks with highest IDR score, and took the 150bp region around each peak summit as positive examples. Negative examples (flanks) were taken 300 bp upstream of positive examples. Overlapping regions and sequences containing Ns are removed. For each TF data set of ChIP-seq peaks and flanks, we made binary predictions based on their similarity to the corresponding TF label in BindSpace and measured performance by F1 score. We then compared with predictions made by FIMO scoring using the ‘Jolma2013’ motifs with a cutoff of *P* < 1e-4.

### ATAC-seq binding versus non-binding evaluation

All ATAC-seq peaks were trimmed to 300bp centered at the peak summit. All TF ChIP-seq peaks were resized to 150bp centered at the peak summit. If an ATAC-seq peak overlaps with a resized TF ChIP-seq peak, we consider that ATAC-seq peak to be bound by the corresponding TF; otherwise, it is unbound by the TF. For this evaluation, we removed any TFs that were bound to < 10% of the ATAC-seq peaks.

### ChIP-seq multiclass evaluation

We resized all ChIP-seq to be 150bp around summit. For each experiment, we took the top 10,000 peaks with most significant IDR and performed multiclass classification of each peak using BindSpace or FIMO motif scoring using TFs in the same family. We evaluated performance by determining the percentage of the top 10,000 peaks that were correctly classified.

## Acknowledgements

We thank Jason Weston for suggesting the StarSpace algorithm and providing access to his code. We also thank William Stafford Noble, Jeffrey Bilmes, and Jacob Schreiber for helpful comments on the project. This work was supported by NIH/NHGRI U01 award HG009395. Supports for H.Y. was also provided by the Tri-Institutional Training Program in Computational Biology and Medicine.

## Reference

1. Berger, M. F. et al. Compact, universal DNA microarrays to comprehensively determine transcription-factor binding site specificities. Nat. Biotechnol. 24, 1429–1435 (2006).

2. Warren, C. L. et al. Defining the sequence-recognition profile of DNA-binding molecules. Proc. Natl. Acad. Sci. 103, 867–872 (2006).

3. Gordân, R. et al. Genomic Regions Flanking E-Box Binding Sites Influence DNA Binding Specificity of bHLH Transcription Factors through DNA Shape. Cell Rep. 3, (2013).

4. Maerkl, S. J. & Quake, S. R. A systems approach to measuring the binding energy landscapes of transcription factors. Science (80-.). 315, 233–237 (2007).

5. Jolma, A. et al. DNA-binding specificities of human transcription factors. Cell 152, 327–339 (2013).

6. Ghandi, M., Lee, D., Mohammad-Noori, M. & Beer, M. A. Enhanced Regulatory Sequence Prediction Using Gapped k-mer Features. PLoS Comput. Biol. 10, (2014).

7. Setty, M. & Leslie, C. S. SeqGL Identifies Context-Dependent Binding Signals in Genome-Wide Regulatory Element Maps. PLOS Comput. Biol. 11, e1004271 (2015).

8. Yang, L. et al. Transcription factor family‐specific DNA shape readout revealed by quantitative specificity models. Mol. Syst. Biol. 13, 1–14 (2017).

9. Shen, N. et al. Divergence in DNA Specificity among Paralogous Transcription Factors Contributes to Their Differential In Vivo Binding. Cell Syst. 6, 470–483.e8 (2018).

10. Wu, L. et al. StarSpace: Embed All The Things! in AAAI(2018).

11. Melvin, I., Weston, J., Noble, W. S. & Leslie, C. Detecting remote evolutionary relationships among proteins by large-scale semantic embedding. PLoS Comput. Biol. 7, (2011).

12. Grant, C. E., Bailey, T. L. & Noble, W. S. FIMO: Scanning for occurrences of a given motif. Bioinformatics 27, 1017–1018 (2011).

13. Barrera, L. A. et al. Survey of variation in human transcription factors reveals prevalent DNA binding changes. Science (80-.). 351, 1450–1454 (2016).

14. Alipanahi, B., Delong, A., Weirauch, M. T. & Frey, B. J. Predicting the sequence specificities of DNA- and RNA-binding proteins by deep learning. Nat Biotechnol 33, 831–838 (2015).

15. Slattery, M. et al. Cofactor binding evokes latent differences in DNA binding specificity between hox proteins. Cell 147, 1270–1282 (2011).

16. Morgulis, A., Gertz, E. M., Schäffer, A. A. & Agarwala, R. A Fast and Symmetric DUST Implementation to Mask Low-Complexity DNA Sequences. J. Comput. Biol. 13, 1028–1040 (2006).

17. Mikolov, T., Corrado, G., Chen, K. & Dean, J. Efficient Estimation of Word Representations in Vector Space. Proc. Int. Conf. Learn. Represent. (ICLR 2013) 1–12 (2013). doi:10.1162/153244303322533223

18. Simon, N., Friedman, J., Hastie, T. & Tibshirani, R. Regularization Paths for Cox’s Proportional Hazards Model via Coordinate Descent. J. Stat. Softw. 39, (2011).

19. Van Der Maaten, L. & Hinton, G. Visualizing Data using t-SNE. J. Mach. Learn. Res. 9, 2579–2605 (2008).

20. van der Maaten, L. Accelerating t-SNE using Tree-Based Algorithms. J. Mach. Learn. Res. 15, 3221–3245 (2014).

21. Krijthe, J. H. {Rtsne}: T-Distributed Stochastic Neighbor Embedding using Barnes-Hut Implementation. (2015).

22. Litzenburger, U. M. et al. Single-cell epigenomic variability reveals functional cancer heterogeneity. Genome Biol. 18, (2017).

23. Buenrostro, J. D., Giresi, P. G., Zaba, L. C., Chang, H. Y. & Greenleaf, W. J. Transposition of native chromatin for fast and sensitive epigenomic profiling of open chromatin, DNA-binding proteins and nucleosome position. Nat. Methods 10, 1213–8 (2013).

